# InteracTor: Feature Engineering and Explainable AI for Profiling Protein Structure-Interaction-Function Relationships

**DOI:** 10.1101/2025.04.10.648139

**Authors:** Jose Cleydson F. Silva, Layla Schuster, Nick Sexson, Melissa Erdem, Ryan Hulk, Matias Kirst, Marcio F. R. Resende, Raquel Dias

## Abstract

Characterizing protein families’ structural and functional diversity is essential for understanding their biological roles. Traditional analyses often focus on primary and secondary structures, which may not fully capture complex protein interactions. Here we introduce InteracTor, a novel toolkit that extracts multimodal features from protein three-dimensional (3D) structures, including interatomic interactions like hydrogen bonds, van der Waals forces, and hydrophobic contacts. By integrating Explainable AI (XAI) techniques, we quantified the importance of the extracted features in the classification of protein structural and functional families. InteracTor’s interpretable features enable mechanistic insights into the determinants of protein structure, function, and dynamics, offering a transparent means to assess their predictive power within machine learning models. Interatomic interaction features extracted by InteracTor demonstrated superior predictive power for protein family classification compared to features based solely on primary or secondary structure, revealing the importance of considering specific tertiary contacts in computational protein analysis. This work provides a robust framework for future studies aiming to enhance the capabilities of models for protein function prediction and drug discovery.

**AUTHOR SUMMARY:** InteracTor is a computational toolkit designed to enhance our understanding of protein structure and function by focusing on three-dimensional (3D) structural interactions. Unlike traditional approaches that primarily rely on sequence or secondary structure data, InteracTor extracts biologically meaningful features such as hydrogen bonds, van der Waals forces, and hydrophobic contacts, which are critical for protein stability and dynamics. By integrating these features into machine learning models alongside explainable AI methods, InteracTor provides interpretable insights into how specific structural interactions influence protein behavior. Our results demonstrate that tertiary structure features significantly improve the accuracy of protein family classification compared to sequence-based methods alone, underscoring the importance of considering 3D interactions in computational protein analyses. The toolkit’s modular design makes it adaptable for diverse applications, including drug discovery and protein engineering. In a broader context, InteracTor bridges the gap between computational biology and practical applications in medicine and biotechnology by offering a transparent and robust framework for analyzing proteins at a molecular level. This work represents a step forward in leveraging structural data to advance predictive modeling and biological discovery.

## INTRODUCTION

In recent decades, high-throughput sequencing techniques have dramatically expanded protein sequence databases. At the same time, advances in cryo-electron microscopy and deep-learning-based computational structure determination methods, including AlphaFold(1) and RoseTTAFold(2), have transformed protein structure elucidation. Consequently, the surge in available sequence and structural data has catalyzed the development of machine- and deep-learning techniques for predictive modeling. This data has been leveraged to address a variety of challenges, such as identifying non-classical secreted proteins(3),(4) predicting binding affinity(5),(6),(7), and engineering proteins for novel functions(8,9).

Central to these algorithms is feature engineering and encoding, aimed at converting protein sequences and physiochemical properties into machine-readable formats. Ideally, this process captures the attributes most relevant to the predictive targets of interest. Sequence-based feature representations are among the most widely utilized, including amino acid composition, chemical property-based features, k-mers, and alignment-based embeddings. These descriptors effectively simplify sequence information and reduce the data dimensionality while still highlighting broader functional characteristics, sequence patterns, and evolutionary relationships. However, sequence-based methods can suffer from high dimensionality and data sparsity and are limited in their ability to capture critical properties influencing protein function. The incorporation of 3D structural data into the suite of available encodings allows predictive models to have a deeper layer of biological context that can give insight into functional dynamics. This has spurred the development of more comprehensive feature extraction platforms such as iFeatureOmega(10), Pfeature(11), and ProFeatX(12) which incorporate both sequence and secondary structure descriptors.

Pfeature utilizes amino acid sequences, employing binary encoding for chemical elements and leveraging PaDEL software for fingerprint generation. ProFeatX focuses on torsional angle bigrams, providing insights into secondary structure. This diverse array of methodologies underscores the complementary nature of these tools, with each excelling in specific bioinformatics applications. However, these tools do not consider the three-dimensional structure of the protein, nor do they consider the interactions between amino acids within the protein’s three-dimensional framework. Tools like iFeature harness molecular structures to calculate Half Sphere Exposure (HSE) and Solvent Accessible Surface (SAS), but still lack support for the extraction of interatomic interactions and other key structural features(10).

While traditional protein sequence descriptors have been fundamental for many predictive modeling types, encoding interaction features remains a significant yet underutilized strategy for capturing the nuances of protein behavior. Although the recent surge in powerful sequence and structural embeddings that significantly improves predictive performance, they often create uninterpretable ’black-box’ models, making it extremely difficult to understand the relationship between the input features and model predictions, even with advanced Explainable AI (XAI) techniques(13). In contrast, InteracTor’s extracted interatomic interaction features are inherently biologically meaningful, allowing for more direct and transparent application of XAI in downstream models. This enables researchers to pinpoint specific interatomic interaction types responsible for observed protein structural and functional properties, offering a clearer path to biological insight.

Here we present InteracTor, a toolkit for the extraction of three types of protein feature encodings: interaction features, physicochemical features, and compositional features. Interaction features include hydrogen bonds, hydrophobic contacts, repulsive interactions, and van der Waals interactions, each encoding unique aspects of molecular dynamics that play an important role in governing protein function. Specifically, hydrogen bonds and hydrophobic contacts are important for stabilizing secondary and tertiary structures(14),(15). Van der Waals interactions influence molecular complementarity, which is crucial for substrate binding, and mediate transient interactions that can facilitate or destabilize protein structures and complexes(16). Physiochemical property features include accessible solvent area, hydrophobicity, and surface tension, which are implicated in protein folding, stability, solubility, and protein-protein interactions. Compositional features include mono-, di-, and tripeptide composition and amino acid side chain chemical property (CPAASC) frequencies(17),(18),(19). These features determine local spatial arrangements (secondary structure) and the overall 3D folded conformation (tertiary structure) of a protein through the formation of alpha helices, beta sheets, loops, and structural motifs.

By leveraging XAI techniques such as Shapley Additive exPlanations (SHAP)(20), we compared the importance of InteracTor’s extracted interatomic interaction features to classic primary and secondary structure features across multiple machine learning model architectures. Our feature sets directly map to biologically meaningful concepts, enabling users to readily interpret results and validate the logic of explainable AI models, in contrast to abstract embedding or principal component vectors. This approach allowed us to pinpoint the most impactful features for characterizing protein families and elucidate the relative contribution of tertiary structural information to predictive performance. Structural and sequence-based features were complementary and provided a more comprehensive representation of the protein. Our integrative feature encoding and selection approach underscores the complexity and richness of proteins, ultimately advancing our ability to characterize proteins for various applications in structural biology, biotechnology and medicine.

## RESULTS

### Multimodal protein profiling

InteracTor computes 11 different interatomic interaction features and 8 distinct CPAASC that are key for characterizing structure and function of proteins (**Table 1** and **Supplementary Table 1**). In addition to interaction and structural features, our toolkit also extracts classic protein sequence compositional features such as k-mer frequencies (**Supplementary Table 2**), resulting in a total of 18,296 multimodal features extracted (**Table 2**). This provides a comprehensive representation of protein structure, function, and sequence characteristics, enabling in-depth analysis of protein properties across various scales of molecular organization. While we demonstrate InteracTor’s utility through protein function family classification, the toolkit is designed for modular adaptation to diverse structural biology tasks—from drug binding analysis to protein engineering—thanks to its interpretable, feature-driven framework.

**Table 1:**
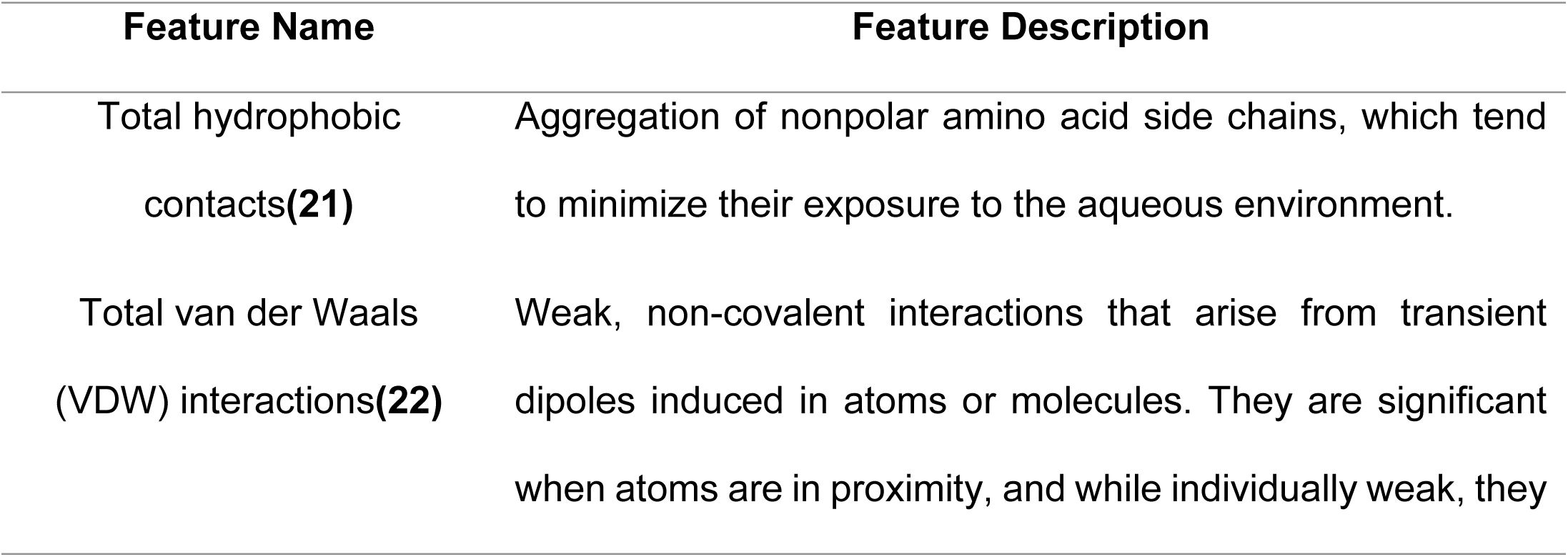
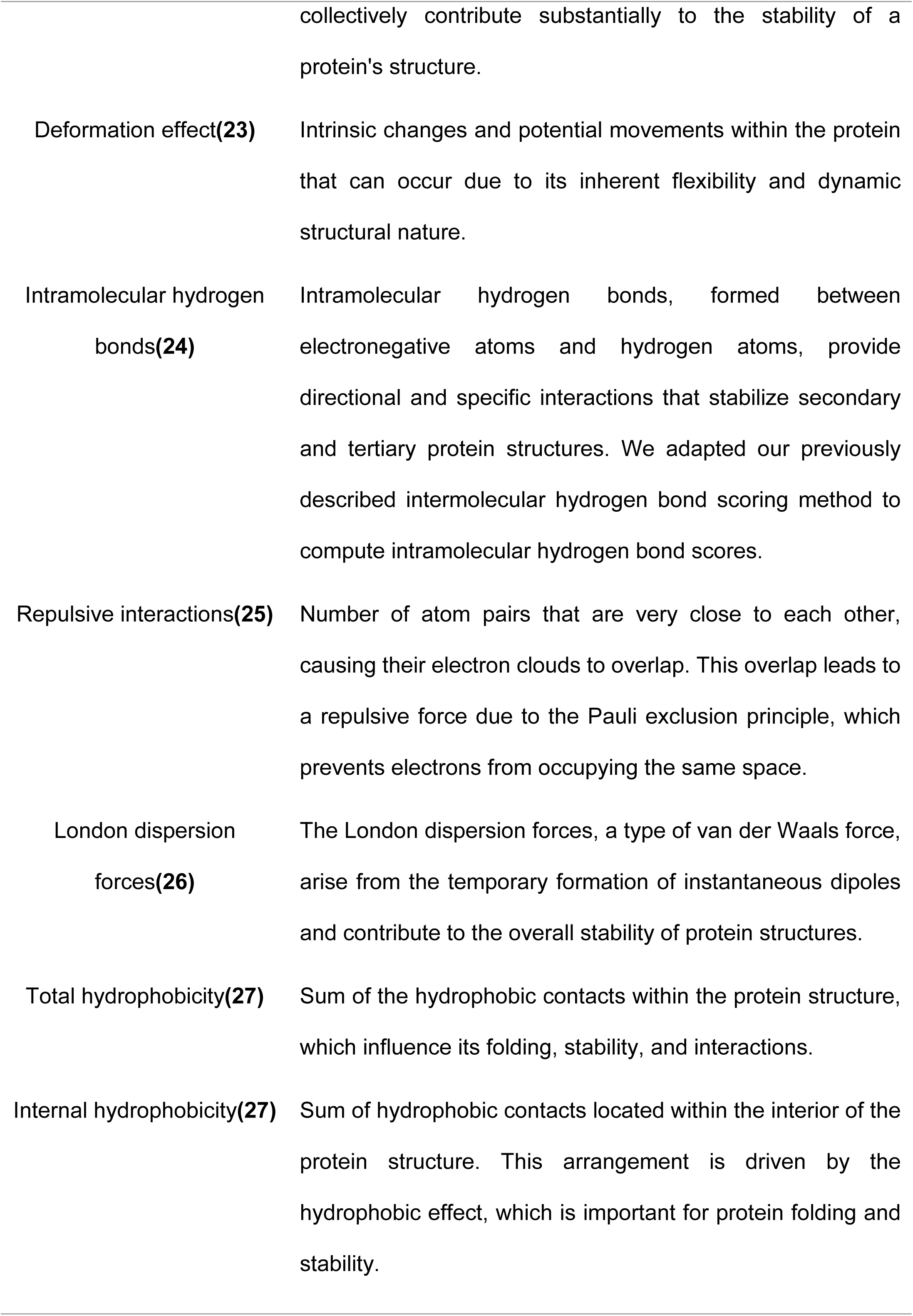
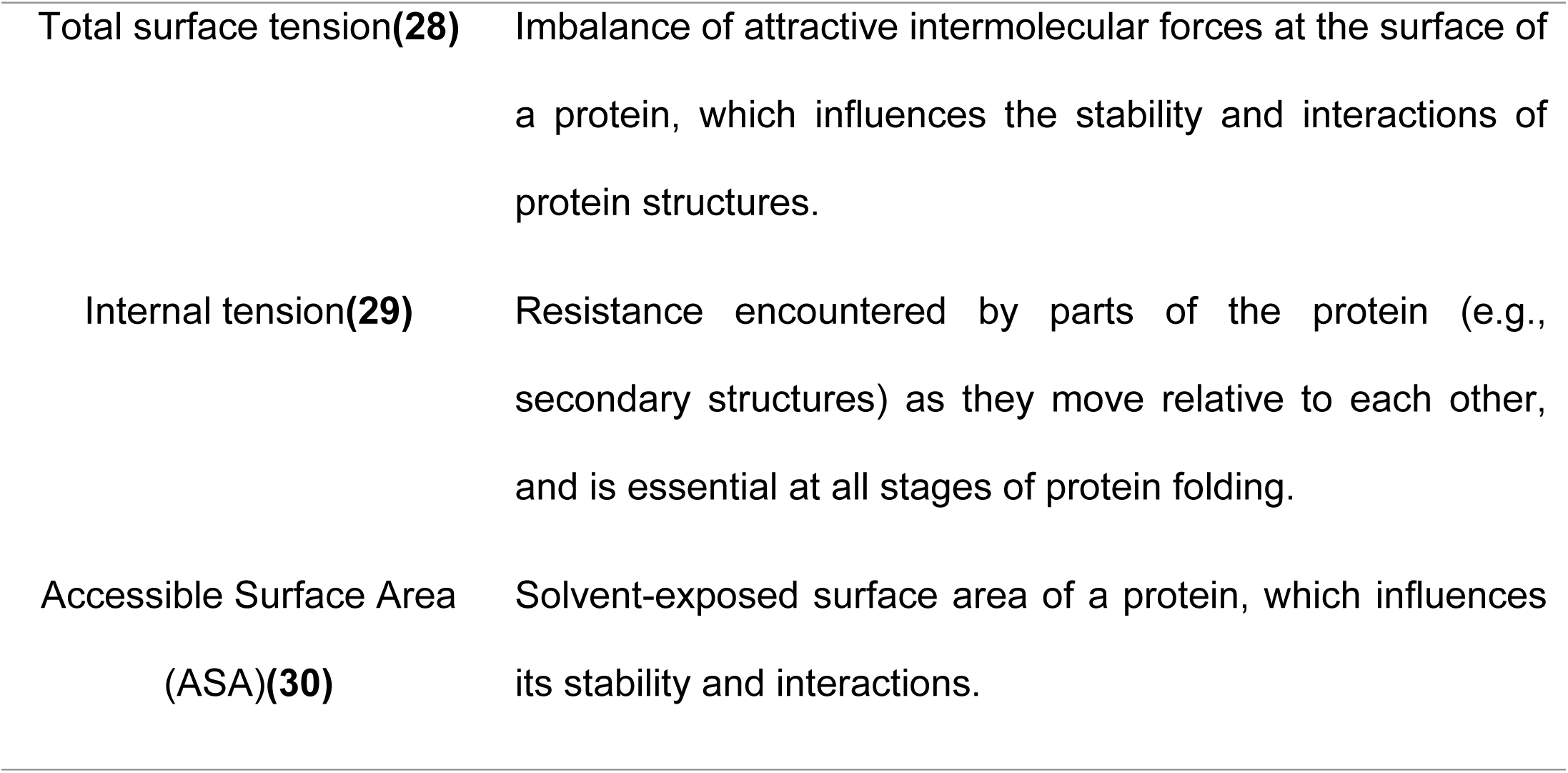
Protein structural features based interatomic interactions.

| Feature Name | Feature Description |
| --- | --- |
| Total hydrophobic contacts(21) | Aggregation of nonpolar amino acid side chains, which tend to minimize their exposure to the aqueous environment. |
| Total van der Waals (VDW) interactions(22) | Weak, non-covalent interactions that arise from transient dipoles induced in atoms or molecules. They are significant when atoms are in proximity, and while individually weak, they |
|  | collectively contribute substantially to the stability of a protein's structure. |
| Deformation effect(23) | Intrinsic changes and potential movements within the protein that can occur due to its inherent flexibility and dynamic structural nature. |
| Intramolecular hydrogen bonds(24) | Intramolecular hydrogen bonds, formed between electronegative atoms and hydrogen atoms, provide directional and specific interactions that stabilize secondary and tertiary protein structures. We adapted our previously described intermolecular hydrogen bond scoring method to compute intramolecular hydrogen bond scores. |
| Repulsive interactions(25) | Number of atom pairs that are very close to each other, causing their electron clouds to overlap. This overlap leads to a repulsive force due to the Pauli exclusion principle, which prevents electrons from occupying the same space. |
| London dispersion forces(26) | The London dispersion forces, a type of van der Waals force, arise from the temporary formation of instantaneous dipoles and contribute to the overall stability of protein structures. |
| Total hydrophobicity(27) | Sum of the hydrophobic contacts within the protein structure, which influence its folding, stability, and interactions. |
| Internal hydrophobicity(27) | Sum of hydrophobic contacts located within the interior of the protein structure. This arrangement is driven by the hydrophobic effect, which is important for protein folding and stability. |
| Total surface tension(28) | Imbalance of attractive intermolecular forces at the surface of a protein, which influences the stability and interactions of protein structures. |
| Internal tension(29) | Resistance encountered by parts of the protein (e.g., secondary structures) as they move relative to each other, and is essential at all stages of protein folding. |
| Accessible Surface Area (ASA)(30) | Solvent-exposed surface area of a protein, which influences its stability and interactions. |

**Table 2:** Overview of protein features extracted.

| Feature type | Number of features | Feature Description |
| --- | --- | --- |
| Interatomic interaction | 11 | Interatomic interactions between residues in a protein, such as hydrogen bonds and hydrophobic contacts. |
| Chemical properties of amino acid side chains (CPAASC) | 8 | The chemical properties of amino acid side chains govern their interactions with other molecules or residues in proteins, encompassing characteristics such as polarity, charge, size, and hydrophobicity. |
| Monopeptide | 26 | Frequency of single residues, or k-mers where k=1. |
| Dipeptide | 675 | Frequency of two residues linked by a peptide bond, or k-mers where k=2. |
| Tripeptide | 17, 576 | Frequency of three residues linked by peptide bonds, or k-mers where k=3. |
| Total | 18,296 |  |

### InteracTor characterizes the variability among distinct protein families

We conducted PCA across all 18,296 features extracted from 20,877 protein structures representing the most abundant protein families and GO terms in the PDB REDO database (**Supplementary Tables 3-4**) to evaluate the overall utility of InteracTor in the characterization of variability among protein families and Gene Ontology (GO) terms. **Fig. 2** shows the first two of 500 selected PCAs (**Supplementary Fig. S1C-D**), which captured 1.72% of the variance across protein families (**Fig. 2B**), and 1.0% of the variance across GO terms (**Fig. 2A**). Except for Peptidase S1 and the Glycosyl hydrolase 5 (cellulase A), the protein families exhibited well-defined clusters in the two-dimensional space (**Fig. 2D**), whereas GO terms exhibited less separation overall (**Fig. 2A**). This difference may be attributed to the inconsistent accuracy of GO annotations across non-model organisms(31).

**Fig. 1.**
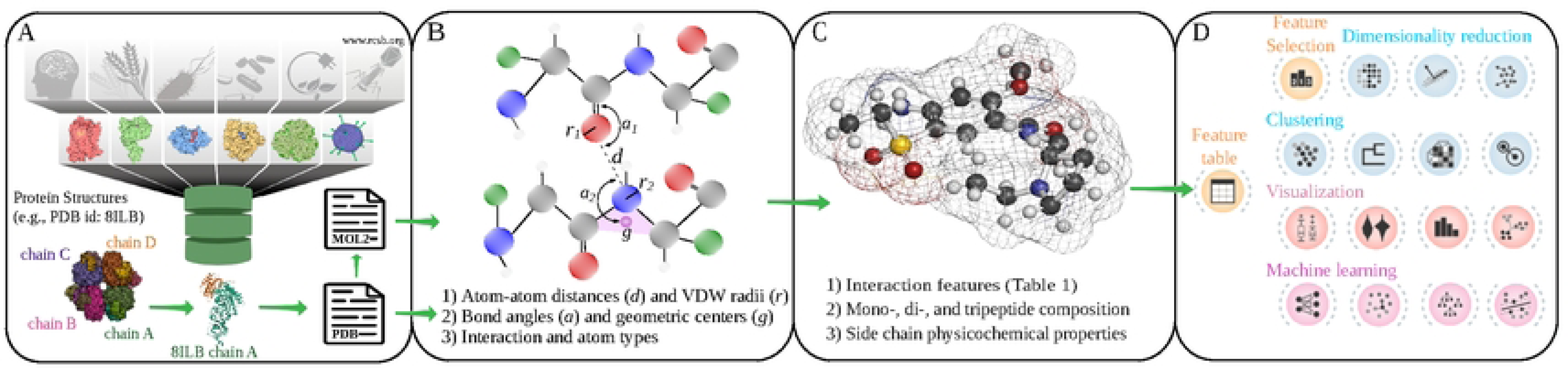
Pipeline of the InteracTor Algorithm. (A) Extensive database of PDB files, where each chain is split into an individual PDB file and converted to MOL2 format. (B) The following metrics are calculated: atom-atom distances (d) and van der Waals radii (r); bond angles (a); geometric centers (g); interaction types and atom types. Molecular representation with carbon atoms in gray, oxygen in red, and nitrogen in blue. (C) The following features are calculated: interatomic interaction features (**Table 1**); side chain physicochemical properties (**Supplementary Table S1**); mono-, di-, and tripeptide composition (**Supplementary Table S1**). (D) The extracted features are exported in tabular format, which then is utilized for feature selection, dimensionality reduction, clustering, visualization, and machine learning.

**Fig. 2.**
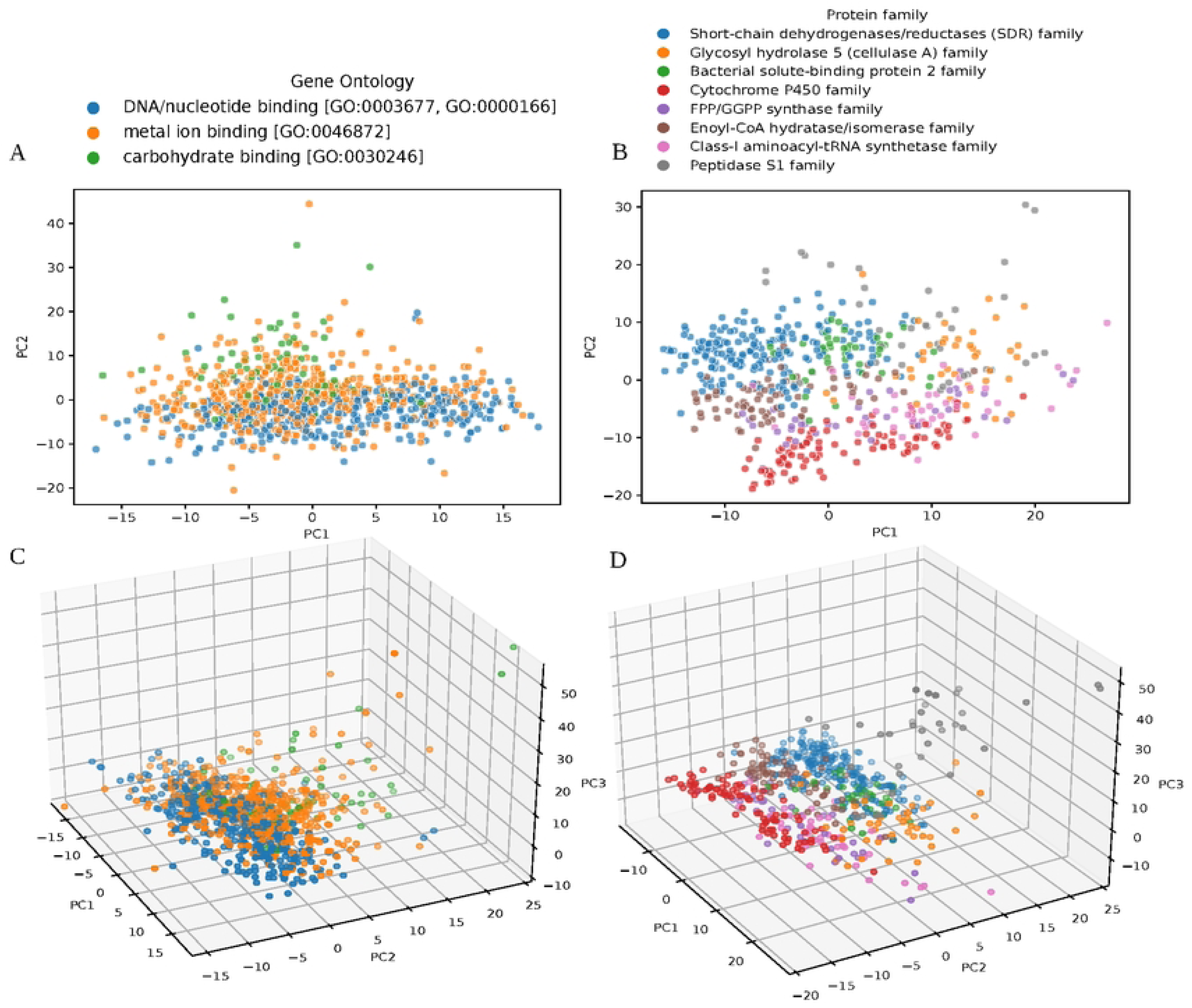
Analysis of Protein Families and Their Molecular Functions. A) Clustering of three representative groups based on molecular functions. B) Clustering of the most prominent protein family, showcasing distinct traits unique to that family. C) Three-dimensional PCA visualization of molecular functions, capturing the variance and complexity within different functional categories. D) Three-dimensional PCA representation of protein families.

Three-dimensional projections provided clearer clustering of the data, allowing a distinct visualization of relationships among the protein families. The first three principal components captured 2.40% of the variance across protein families (**Fig. 2C, Fig. S1A**), as well as 1.38% of the variance across GO terms (**Fig. 2D, Fig. S1B**). The 3D visualization also improved the separation of Peptidase S1 and Glycosyl hydrolase 5 families, as well as the separation of GO groups, allowing for a more nuanced interpretation of functional relationships among groups (**Fig. 2C**).

### Mutual Information scoring identifies key features across protein families

To enhance clustering quality and classification accuracy in downstream analyses, InteracTor performs feature selection using Mutual Information (MI) scoring. All features were ranked by MI score to prioritize those most relevant to the target variables. **Fig. 3A** shows the distribution of MI scores used for feature selection. The distribution exhibits a primary mode, or large peak, which corresponds to features with low MI scores (MI<0.2), reflecting background noise and less informative features. The remaining lower peaks (MI>=0.2) represent a subset of 354 features with informational content effectively distinguished from the background noise peak. Among the top 100 high MI scoring features are 9 interatomic interactions, 2 CPAASC features, and 89 sequence composition features (**Supplementary Table 5**). Among the 12 most highly ranked features across protein families (**Fig. 3B**) are hydrogen bonds (MI=0.775), total surface tension (MI=0.763), London dispersion forces (MI=0.758), repulsive interactions (MI=0.722), internal tension (MI=0.708), Accessible Surface Area (ASA) (MI=0.694), hydrophobic contacts (MI=0.561), TG frequency (MI=0.562), internal hydrophobicity (MI=0.561), VN frequency (MI=0.556), total hydrophobicity (MI=0.539), and GG frequency (MI=0.509).

**Fig. 3.**
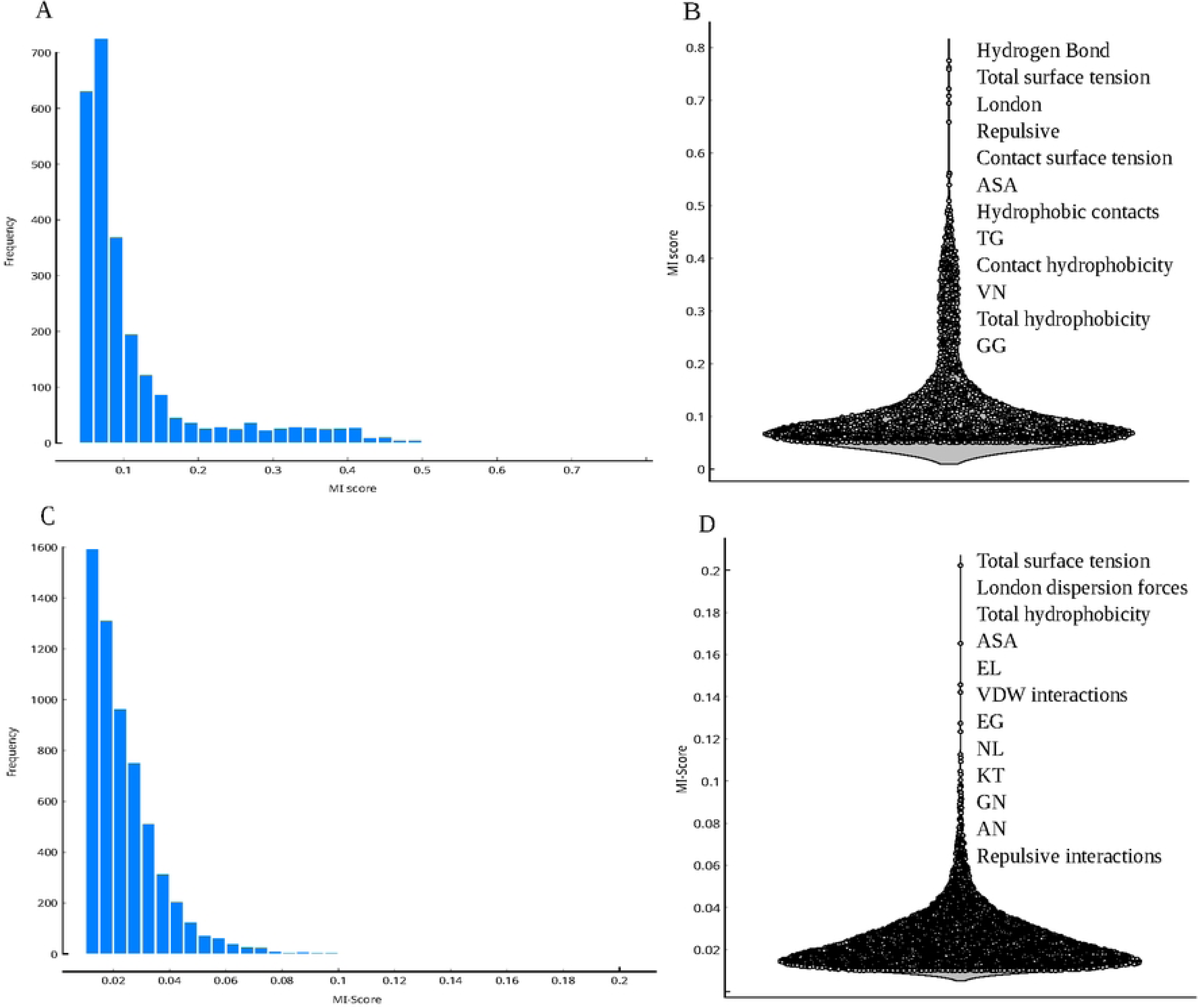
Mutual Information (MI) Scores Across Protein Families and Molecular Functions. A) Frequency distribution of MI scores for each protein family B) Density distribution of MI scores across protein families, focusing on the top 12 ranked features. C) Frequency distribution of MI scores for molecular functions. D) Density distribution of MI scores for molecular functions, focusing on the top 12 ranked features.

For GO terms, the MI score analysis revealed values ranging from approximately 0.06 to 0.1, as shown in **Fig. 3C**. While several peaks are well-defined, indicating key features, the MI scores are notably lower compared to those for protein families. **Fig. 3D** presents a violin plot illustrating the distribution of MI scores, with a high density of values between 0.1 and 0.2. High MI scoring features include total surface tension (MI=0.202), London dispersion forces (MI=0.165), total hydrophobicity (MI=0.146), ASA (MI=0.142), EL frequence (MI=0.127), VDW interactions (MI=0.123), EG, NL, KT, GN, AN, (MI=0.113, 0.111, 0.109, 0.105, and 0.103, respectively) and repulsive interactions (MI=0.100). Despite the lower MI scores, these features show a significant association with protein families and GO categories (Wilcoxon test p≤2.2x10^-16^).

### Clustering of selected features reveals distinct patterns and relationships among protein families

Hierarchical clustering grouped the dataset into 19 distinct clusters across eight families (**Fig. 4A**), revealing complex internal organization. The Short-chain dehydrogenases/reductases (SDR) family exhibited the most extensive dispersion, spanning seven clusters (C1, C6, C10, C11, C12, and C13), highlighting its functional diversity. Three distinct clusters (C17, C19, C20) encompassed the Cytochrome P450 family. Similarly, the Peptidase S1 family was distributed across three clusters (C8, C9, C16), suggesting potential functional specialization within this protease group. The Enoyl-CoA hydratase/isomerase family was found in two clusters (C2, C3), indicating possible sub-functionalization. In contrast, several protein families demonstrated a more focused distribution, each confined to a single cluster: Bacterial solute-binding protein 2 (C4), FPP/GGPP synthase (C14), Glycosyl hydrolase 5 (cellulase A) (C15), and Class-I aminoacyl-tRNA synthetase (C18).

**Fig. 4.**
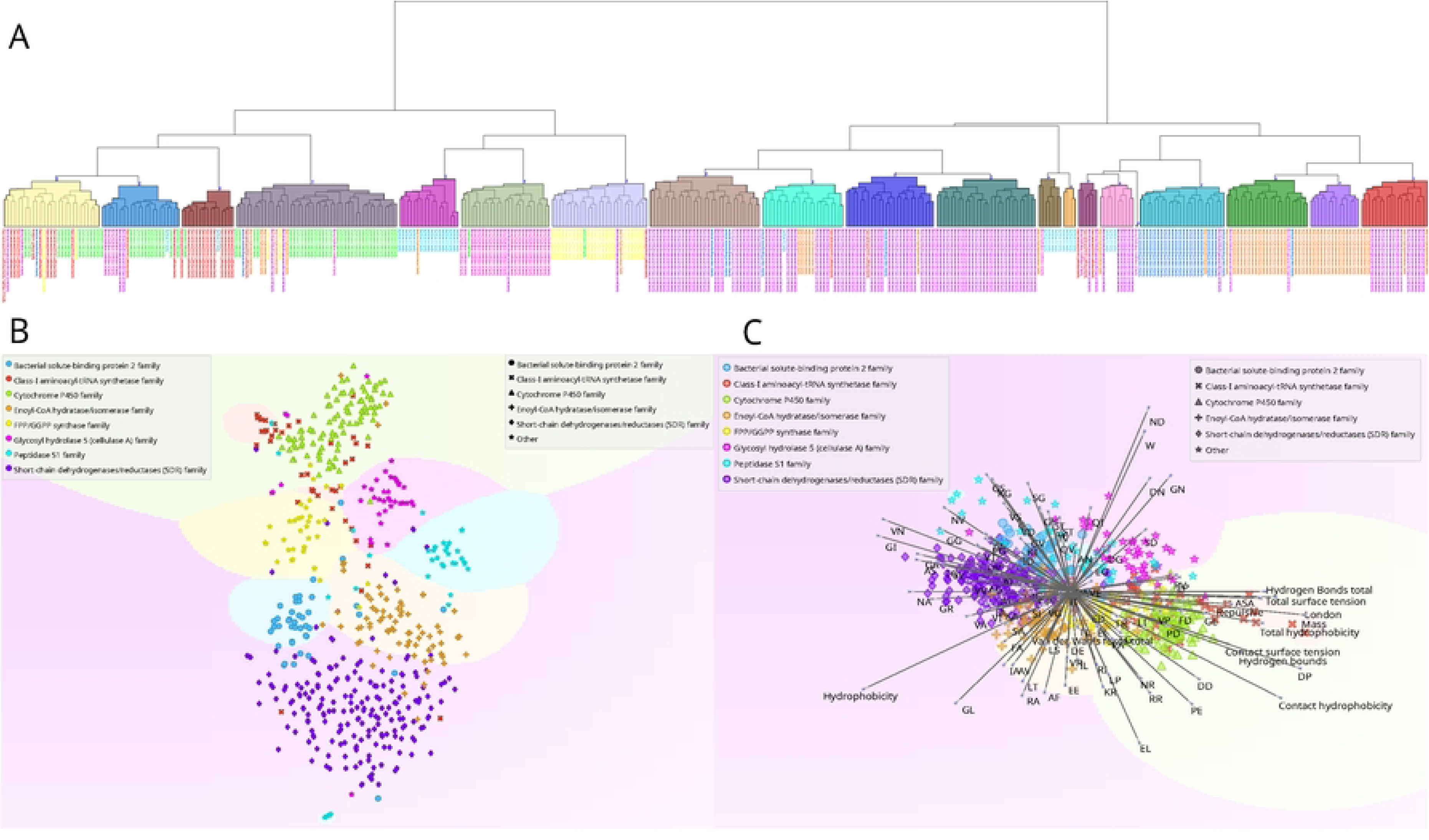
Clustering and Visualization of Protein Families. A) Hierarchical clustering of protein families, depicting relationships and groupings based on structural and functional similarities. B) t-SNE plot depicting inter-family relationships within a reduced-dimensional space. C) Optimized projection plot highlighting the most significant features for each group, illustrating key attributes that differentiate protein families.

Complementary t-SNE clustering analysis identified 9 groups (**Fig. 4B**), not only corroborating these groupings but also unveiling finer details of inter-group relationships. While hierarchical clustering effectively segmented proteins with similar functions, t-SNE provided a more nuanced separation. Occasional cross-family similarities were highlighted with some datapoints from different families appearing in unexpected clusters. Multivariate visualization further supported the role of interatomic interaction features (**Fig. 4C**), which were predominantly observed in the Cytochrome P450 and Glycosyl Hydrolase 5 (cellulase A) families (Wilcoxon p≤4.9x10^-4^, **Supplementary Table 6**). In contrast, dipeptide features were more prevalent among other protein families, indicating a varied functional landscape shaped by distinct sequence composition patterns. (Wilcoxon p≤0.031, See **Supplementary Table 6**).

### InteracTor’s feature selection effectively reduces dimensionality of models without compromising model performance

Across all feature sets described in **Table 2**, ensemble models, especially Histogram Gradient Boosting and Random Forest, consistently ranked among the top performers in protein family classification, achieving 0.71-0.80 F1-score on the test dataset after 80%-20% train-test split, indicating that the complex relationships captured by InteracTor’s features are effectively leveraged by these models’ ability to combine multiple decision trees (**Fig. 5A**). This trend was consistent across different accuracy metrics such as accuracy, precision, recall, and MCC **(Supplementary Tables 7A** and **7B)**. CPAASC was the feature type that showed the lowest accuracy across all feature subsets (Tukey’s post hoc test p≤0.05, **Supplementary Table 7A, 8** and **Fig.5B**). MI score-based feature selection and further F1 score comparison across models showed that reducing InteracTor’s feature set to the top 100-500 features achieved comparable performance to using all 18,296 features (Tukey’s post hoc test p≥0.82). The smallest feature set selected via MI score (top 100) showed similar performance relative to larger subsets as well (Tukey’s post hoc test p≥0.12 versus top 200-500, **Supplementary Table 8**). These results highlight the effectiveness of our feature selection approach for optimizing model performance and interpretability.

**Fig. 5.**
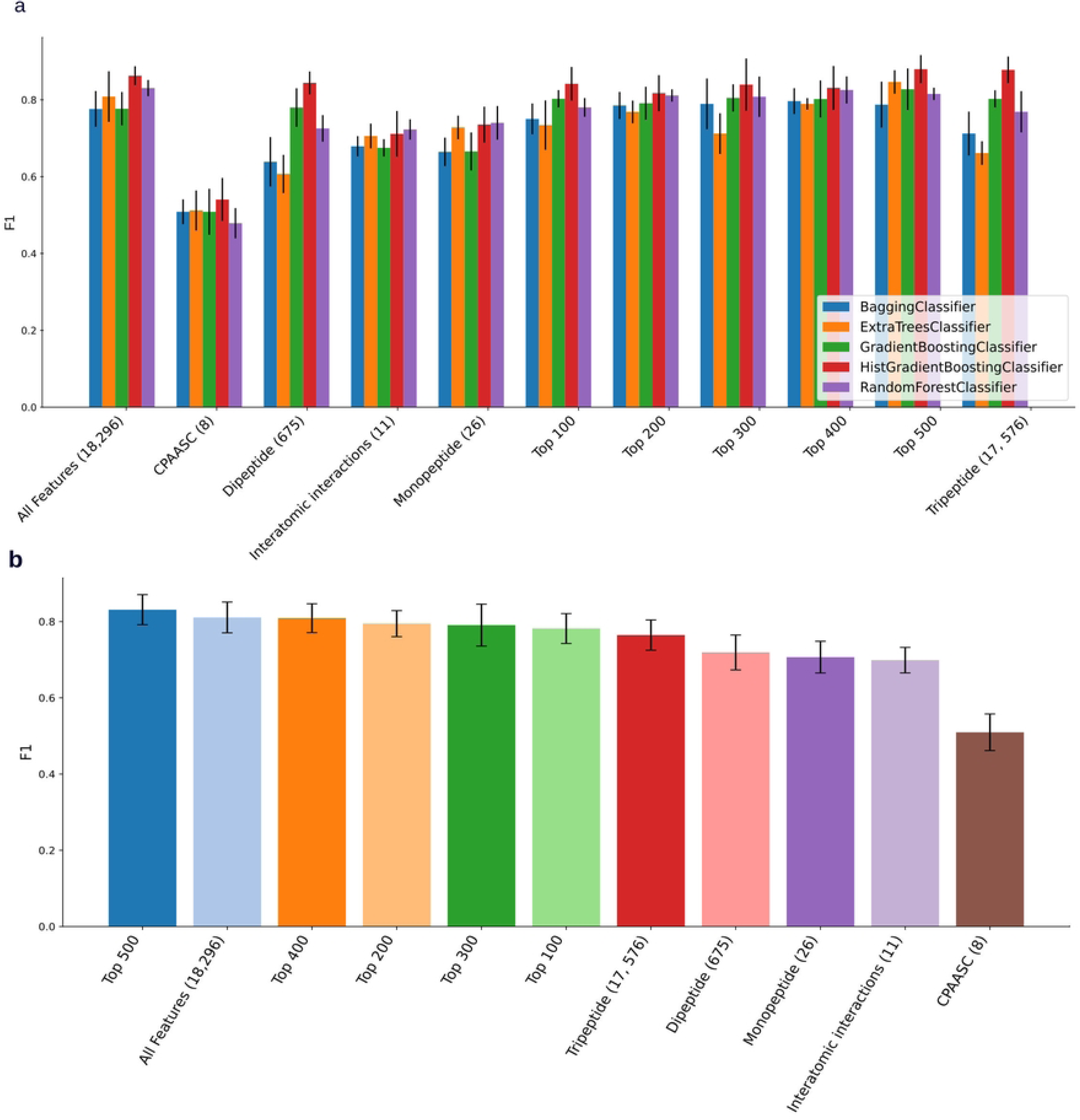
Machine learning benchmark across different feature sets. A) F1 scores were computed for several classification models (shown as colored bars) across different feature sets. Feature sets were ranked by their impact on model performance, with higher F1 scores indicating superior predictive accuracy. B) Performance of the best model (Histogram Gradient Boosting) across different feature sets, showing mean validation accuracy after 5 runs.

### XAI reveals key features for protein family prediction models

We employed eXplainable Artificial Intelligence (XAI) techniques, specifically the SHapley Additive exPlanations (SHAP) method(20), to analyze the impact of InteracTor’s features on machine learning models for protein family prediction. The top 20 features showed SHAP values ranging from 0.25 to 1.75 and showed heterogenous contributions to the models’ predictions across different protein families (**Fig. 6, Supplementary Fig. S2**). The most important features included interatomic interactions and physicochemical properties, such as surface tension and hydrophobicity, as well as specific atomic interactions like repulsive interactions and hydrogen bonds. Additionally, certain peptide composition patterns, such as those involving amino acids P, CLG, PP, H, and W, were also among the most important features. These findings indicate that a combination of multimodal feature types, including structural properties, interatomic interactions, and amino acid composition, critically affects the model’s capacity to differentiate protein families.

**Fig. 6.**
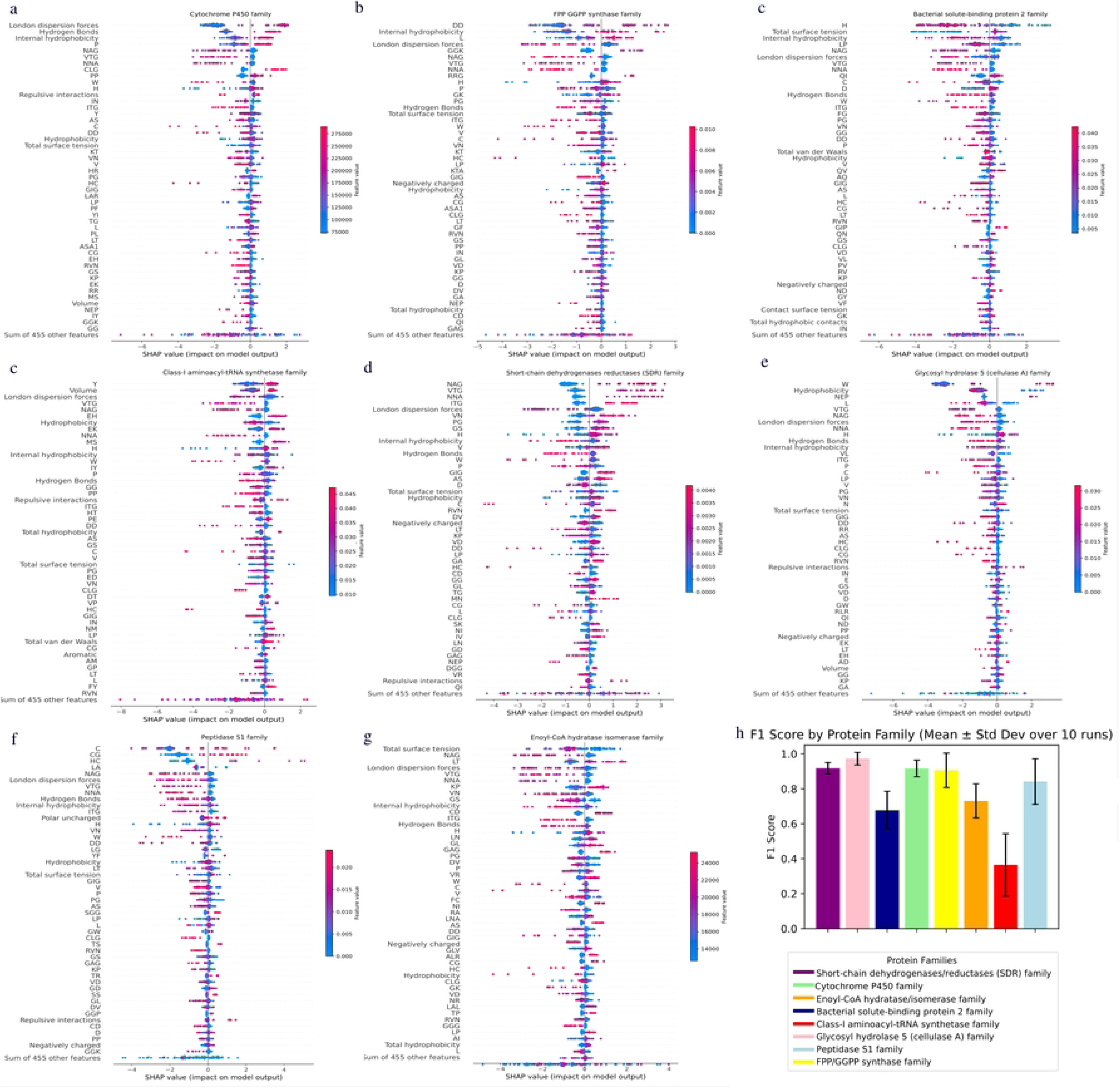
Feature Importance (SHAP) in Protein Family Classification. A-G) The plot ranks features by their impact on model predictions, illustrating contribution distributions. Each dot represents a protein, with colors indicating feature values: red for high, blue for low. The horizontal spread of dots reflects the density of examples at given feature values. Negative SHAP values (left) indicate a reduced likelihood of classification, while positive SHAP values (right) suggest an increased likelihood. SHAP values were computed using family-specific explicands and baselines, ensuring consistent cross-group comparisons. h) Classification model performance measured by the F1 Score for each protein family. The bars represent the mean F1 Score obtained from 10 independent runs, with error bars corresponding to the standard deviation.

The features were ranked differently by SHAP values across different protein families, with each protein family type exhibiting a unique association pattern. For instance, London dispersion forces, hydrogen bonds, and internal hydrophobicity had a positive impact on predictions for the Cytochrome P450 family (**Fig. 6A, Supplementary Fig. S2A**). However, these same features showed negative impacts for other families, including Bacterial solute-binding protein 2, Class-I aminoacyl-tRNA synthetases, and Short-chain dehydrogenases reductases (**Fig. 6B-G, Supplementary Fig. S2B-G**). FPP/GGPP synthases also exhibited a distinct pattern and ranking of SHAP values relative to other families, with key contributors from dipeptide DD, internal hydrophobicity, london dispersion forces, and several specific amino acid combinations including tripeptide RRG and monopeptide H (**Fig. 6B, Supplementary Fig. S2B**).

## DISCUSSION

### A novel algorithm for improved characterization of protein structural properties

Our protein feature extraction algorithm distinguishes itself by focusing on the tertiary structure of proteins, unlike other approaches that rely on primary and secondary structures(10,32,33). By analyzing three-dimensional relationships, the algorithm captures intramolecular interactions that underpin protein structure and function. Extracting features directly from the tertiary structure allows the algorithm to uncover additional patterns in the chemical properties of amino acids and their interactions, offering a comprehensive understanding of the structural and functional dynamics of proteins(34). This enhanced perspective provides new insights into the complex mechanisms governing protein behavior, facilitating advancements in protein engineering and drug design(35). The graphical representations (**Fig. 3-6**) provide a comprehensive overview of protein characteristics as well, assisting in the visualization and interpretation of complex structural and physicochemical properties across protein families.

The feature selection process, guided by MI scores, was crucial in identifying the most effective features for distinguishing protein families (**Fig. 3**). Prominent peaks in MI scores indicated that features such as hydrogen bonds, total surface tension, and contact hydrophobicity are particularly influential in differentiating protein families. This prioritization of key features enhances our ability to effectively distinguish between various protein structural and functional groups. The identification of 9 out of 11 interatomic interactions features among the top 12 features with the highest MI scores underscores their importance in differentiating protein families. This finding defines the importance of specific molecular interactions, such as hydrogen bonds and hydrophobic contacts, in defining protein structure and function, providing valuable insights into the fundamental principles governing protein family diversity.

**Fig. 4** shows clear separations of specific clusters, highlighting the effectiveness of the approach in identifying grouping patterns among protein families. The groupings not only reflected complex interatomic relationships between different features but also demonstrated the formation of distinct clusters within dipeptide feature groups. The Peptidase S1 family, Glycosyl hydrolase 5 (cellulase A) family and Short-chain dehydrogenases/reductases (SDR) family demonstrated clustering of dipeptide feature groups. This result confirms the method’s ability to highlight relevant functional patterns and distinguish between different protein family categories based on the observed interatomic interactions.

### Clustering reveals complex structure and functional diversity in protein families

Hierarchical clustering effectively delineated differences among protein families such as short-chain dehydrogenases/reductases (SDR), Cytochrome P450, Enoyl-CoA hydratase/isomerase, Bacterial solute-binding protein 2, Class-I aminoacyl-tRNA synthetase, Glycosyl hydrolase 5 (cellulase A), Peptidase S1, and FPP/GGPP synthase. Complementary analysis with t-SNE allowed the detection of subtle similarities and differences among the protein families and offering deeper insights into their functional connections (**Fig. 4**, **Supplementary Table 6**). Multivariate visualization allowed dynamic adjustments to identify unique aspects of protein families, including specific features contributing to the clustering process, yielding valuable insights into the functional diversity within and between protein families. This integrative approach led to a deeper understanding of the complex relationships and functional diversity inherent across protein families, revealing details on the interplay of intermolecular interactions, physicochemical properties, and composition of proteins. This holistic methodology not only enhances our comprehension of protein family dynamics but also establishes a robust framework for future research. The extracted features have the potential to directly impact the performance of machine learning and deep learning models by offering a new feature engineering and feature selection tool. By providing new insights on protein structural and functional properties, our approach supports advancements in structural biology, synthetic biology and in the characterization proteins of unknown function (PUFs)(36–39). Additionally, these features can contribute to emerging fields such as quantum biology(40–44), facilitating a deeper understanding of biological roles and interactions across diverse protein families.

### XAI reveals key features for classification of protein function families

Our XAI results show that a combination of physicochemical properties, atomic interactions, and peptide motifs enhances protein family classification accuracy. This approach underscores the importance of integrating protein structural, physicochemical and sequence features in computational modeling, assembling groundwork for advanced protein analysis methods. By integrating these features, we open a new pathway for understanding protein structure, activity, and function. Feature selection and XAI analysis revealed that atomic-level interactions, specifically hydrogen bonds and London dispersion forces, in conjunction with distinct peptide composition patterns (e.g., proline, cysteine-leucine-glycine, and diproline sequences) were pivotal for identifying protein families. Moreover, the application of XAI methodologies elucidated the differential impact of these features across various protein families (**Fig. 6**).

The heterogeneity in feature importance can be attributed to the structural and functional diversity of the features among protein families. For instance, the importance of London dispersion forces(45) and internal hydrophobicity(46) in Cytochrome P450 classification emphasizes the crucial role these interactions play in maintaining its tertiary structure. Conversely, the inverse correlation of these same features in families such as Peptidase S1 indicates that alternative factors such as hydrophobicity, the mono-dipeptide, and the notable presence of specific catalytic domains may exert a more substantial influence on classification outcomes(47) (**Supplementary Fig. S2F**).

Our findings align with previous studies that emphasize the significance of atomic interactions and physicochemical properties in protein classification(5,10–12,17–19). Other similar features based in sequence property have been widely reviewed(48,49). Our proposed set of interatomic interaction features were often among the top ranked features contributing to the accurate classification of protein families, complementing other feature types (**Supplementary Tables 5-8**). By leveraging XAI, we successfully quantified and interpreted the specific impact of each feature- an aspect that conventional approaches could not comprehensively address-offering deeper insights into the biological mechanisms underlying the model’s performance.

The capacity to identify key features for protein classification has implications for drug design and protein engineering because understanding molecular interactions is decisive. InteracTor enables interpretable modeling tasks such as protein feature engineering, feature selection, and XAI, supporting applications like drug discovery (e.g., binding site analysis), protein function prediction, and structure-guided protein design. InteracTor applies XAI principles—such as feature importance attribution and structural interpretability—to map interatomic interactions to functional outcomes. This bridges the gap between black-box protein models and actionable insights, enabling targeted protein engineering (e.g., optimizing binding affinity or stability by prioritizing CPAASC features) and error diagnosis (e.g., flagging repulsive VDW clashes in misfolded designs).

### Mechanistic insights on protein structure, activity, and function

The features extracted by InteracTor provide critical insights into the structural and functional diversity of analyzed protein families. Hydrogen bonds were among the top predictors for classifying SDR proteins. SDR proteins have a conserved α/β folding pattern with a central beta sheet flanked by 2-3 α-helices from each side, forming a classical Rossmann-fold motif for nucleotide binding. Hydrogen bonds are known to be essential in the stabilization of the Rossmann fold and the NAD(P)(H)-binding region of SDRs, which is consistent with our findings(50),(51). ASA and total hydrophobic contacts were key features in the classification of Cytochrome P450 family. The importance of hydrophobic contacts in Cytochrome P450 function has been highlighted by studies showing that hydrophobic residues are pivotal in complex formation with their redox partners (52). Cytochrome P450s also have deeply buried active sites that are connected to the solvent by a network of channels exiting at the distal surface of the protein, which could be reflected by the high contribution of ASA in the classification of this protein family(53).

The Bacterial solute-binding protein 2 family displayed a balance of repulsive interactions and internal tension, likely linked to their dynamic conformational states required for substrate transport(54). Proteins from the Class-I aminoacyl-tRNA synthetase family were characterized by high deformation effect, supporting their complex multi-step role in amino acid activation and tRNA binding(55). The Glycosyl hydrolase 5 (cellulase A) family exhibited pronounced hydrophobic contacts and surface tension, aligning with their function in breaking down robust carbohydrate polymers(56). Hydrophobicity-related features were key predictors for the Peptidase S1 family. Chymotrypsin-like enzymes within the S1 family have a hydrophobic S1 pocket, which allows them to cleave peptide bonds following medium to large hydrophobic amino acids such as tyrosine, phenylalanine, and tryptophan, which is consistent with our findings(57). The Peptidase S1 family also contains a catalytic site that is typically preceded by a block of hydrophobic residues as well, which is also consistent with our results(58).

Lastly, the FPP/GGPP synthase family demonstrated a notable deformation effect, which is consistent with their known versatility by accommodating diverse substrates during isoprenoid biosynthesis(59) These observations highlight InteracTor’s utility in characterizing protein families based on their global structural features, offering a foundation for future studies exploring interaction-specific mechanisms within these families.

## RESOURCE AVAILABILITY

### Data and code availability

InteracTor is available as an open-source Python package on <u>GitHub</u> (https://github.com/Dias-Lab/InteracTor), providing a wealth of details in both code and documentation. The repository includes comprehensive implementation details, example datasets, and usage guidelines to ensure transparency and reproducibility. Users can explore, modify, and contribute to enhance its capabilities.

## ACKNOWLEDGEMENTS

The authors gratefully acknowledge UF Research Computing for providing computational resources and support that have contributed to the research reported in this publication (http://www.rc.ufl.edu).

This work was also supported by UF LIFT AI funding of University of Florida’s Institute of Food and Agricultural Sciences (UF/IFAS).

## AUTHOR CONTRIBUTIONS

R.D. designed and supervised the study and contributed to manuscript writing. M.F.R.R and M.K. contributed to manuscript writing. J.C.F.S. developed the algorithm, built the computational tools, performed the analysis, and wrote the manuscript. L.S and N.S. were responsible for the data processing and contributed to manuscript writing. M.E. and R.H. contributed to software testing and manuscript writing. All authors read and approved the final manuscript.

## METHODS

### Algorithm overview

InteracTor algorithm consists of a sequence of steps for analyzing protein structures and calculating various features from proteins’ interatomic interactions, physicochemical properties of residues, and peptide composition (**Fig. 1**). The following is a description of each step:

*<u>Step 1</u>.* Extract atom, residue, and sequence information from PDB file (**Fig. 1A**): this process involves parsing the Protein Data Bank (PDB) file to obtain the atomic types, 3D coordinates, and the amino acid sequence of the protein.

*<u>Step 2</u>.* Extract additional atomic information from MOL2 file (**Fig. 1A**), including covalent bond mapping and types (e.g., single, double, triple bonds, aromatic rings, etc.) and sp^2^ hybridization.

*<u>Step 3.</u>* Calculate atom-atom distances **(Fig. 1B**): the algorithm computes van der Waals (VDW) radii and distances between pairs of atoms not covalently bound within the protein structure, which is used to identify potential interatomic interactions within the protein structure. A distance threshold, as described by Dias et al.,(5) is applied to determine whether atoms are interacting.

*<u>Step 4</u>.* Calculate geometric centers (**Fig. 1B**): additional properties are extracted from atoms selected as potentially participating in non-covalent interactions (step 3) and covalently bound to multiple atoms (step 2). This includes the calculation of geometric centers between the selected atoms and their respective covalently bound atoms. These centers are utilized to calculate angles between hydrogen bond donor and acceptor atoms (step 5).

*<u>Step 5.</u>* Calculate angles (**Fig. 1B**): the algorithm then computes the angles between atoms or groups of atoms in the protein structure, which is used to evaluate hydrogen bonding geometry.

*<u>Step 6</u>*. Compute interatomic interaction features and protein physicochemical properties described in **Table 1** and **Fig. 1C**.

*<u>Step 7.</u>* Extract compositional features (**Supplementary Table 2** and **Fig. 1C**): InteracTor calculates the frequencies of single amino acids (monopeptide), pairs of amino acids (dipeptide), and triplets of amino acids (tripeptide) in the protein sequence.

*<u>Step 8</u>*. Extract CPAASC frequencies (**Supplementary Table 1** and **Fig. 1C**): the algorithm computes the frequencies of several physicochemical properties associated with the side chains of the amino acids in the protein sequence.

*<u>Step 9.</u>* Write results and postprocessing (**Fig. 1D**): the final step is to write the features computed in steps 6, 7 and 8 to an output file for further analysis or use in other applications. The toolkit also includes example scripts for downstream analyses such as feature selection via MI scoring, dimensionality reduction via PCA, t-SNE, UMAP, hierarchical clustering and visualization using heatmaps, and protein function classification using machine learning and XAI.

Our toolkit extracts and encodes protein structural features based on physico-chemical properties, amino acids composition, and interatomic interactions. By building upon our previous methods for predicting protein-ligand binding affinity(5), we modified the algorithm to analyze residue-residue interactions within protein structures (**Table 1**). We also included the calculation of CPAASC frequencies(17), (**Supplementary Table 1**), monopeptide(18), dipeptide, and tripeptide composition(19) (**Supplementary Table 2**). In addition to the 20 classic amino acids, our algorithm supports rare residues found in distinct biological systems, including selenocysteine, pyrrolysine, and N-methylvaline. These amino acids, although rare, are key in specific biological processes(60),(61).

### Structural datasets and preprocessing

*Dataset for clustering analyses.* We used 20,877 protein structures from the PDB-REDO database(62), having eliminated redundant (50% sequence similarity) and small proteins (<50 residues) using Biopython and in-house scripts (see Data and Code Availability). PDB-REDO was used to enhance structural data quality, improving machine learning model reliability by minimizing errors and inconsistencies (e.g., low resolution, missing atoms, etc.). This approach aligns with research showing that prioritizing data quality over quantity leads to better prediction performance and model robustness(62). Including distant protein relatives with lower identity cutoffs in the study can provide valuable insights into functional conservation and evolutionary relationships as proteins with low sequence identity can still share similar tertiary structures and functions(62). PDB files were converted to MOL2 format using Open Babel(63). We then applied the InteracTor algorithm to extract features from both PDB and MOL2 files.

*Datasets for machine learning and XAI.* We further filtered out proteins with more than 50% sequence identity to minimize data leakage in our machine learning and XAI experiments. We also filtered out structures with low resolution (>2.5 Å) and less than 100 residues in order to reduce noise and potential outliers.

*Datasets selected for demonstration.* We utilized UniProt’s application programming interface (API)(64) to extract Gene Ontology(65) (GO) terms and protein family names based on the PDB accession number(66) to facilitate analysis and demonstration of use cases. We performed power analysis to determine the minimum sample size needed per protein family and GO term for further demonstration of use cases (**Table 1, 2**). For the demonstration of use cases, we selected protein family categories with at least 30 representatives in our dataset to ensure sufficient statistical power for downstream analyses. Similarly, we applied a minimum threshold of 90 annotations for Gene Ontology (GO) terms but further restricted selection to terms directly associated with the binding mechanisms of proteins (e.g., ligand type) to further assess the potential of InteracTor for profiling of protein functions directly associated with ligand or substrate binding. By selecting protein families and GO terms with enough sample sizes, we ensure that our use cases effectively demonstrate our approach’s capabilities. Mapping and power analysis scripts are available in GitHub as well (see Data and Code Availability).

### Feature selection and clustering analysis

We applied MI scoring to quantify the relevance of each feature in distinguishing between different protein families and GO categories(67). We ranked the features by their respective MI scores and selected the top 100 most informative features for further analyses.

We applied PCA to reduce the dimensionality of the data generated by our toolkit and evaluated how the primary components capture the variability among protein families and GO terms(68). In addition to PCA, we also applied t-SNE(69,70) and Freeviz(71) to reduce dimensionality while preserving nonlinear relationships and local structures in data. We used violin plots to visualize the distribution and density of the extracted features across different protein families and GO terms, providing insights into the distribution and central tendencies of the data.

We performed hierarchical clustering using Pearson correlation as the distance metric and the complete linkage method for cluster formation(70). This approach allowed us to identify and visualize the hierarchical relationships between protein families based on similarities among the features extracted by our toolkit. The dimensionality reduction and clustering results were visualized with FreeViz.

### Machine learning benchmark

We generated 11 feature sets to assess their contribution to improving machine learning performance in predicting structural properties. Five datasets comprised individual feature types: interatomic interactions, CPAASC, monopeptides, dipeptides, and tripeptides (**Table 2**). Another five datasets were created using the top 100, 200, 300, 400, and 500 features ranked by MI scores, incorporating a mixture of the five feature types (**Supplementary Table 5**). The eleventh dataset integrated all features extracted by InteracTor (**Table 2**). This approach allowed us to evaluate the relative importance of different feature combinations in enhancing predictive accuracy.

We tested Machine Learning (ML) models to assess the performance of InteracTor’s features in the prediction of protein families. In this experiment, we used 43 ML algorithms implemented in the Python library Scikit-learn (**Supplementary Table 7D**)(72). To evaluate model performance, we randomly shuffled the data, performed 80%-20% train-test split and measured multiclass classification accuracy, precision, F1 score, Mathew’s correlation coefficient (MCC), and precision for each combination of feature set and algorithm. This process was repeated 5 times. We also performed One-Vs-Rest-Classifier cross validation on the best model and feature set to compute F1 scores for each protein family class(70).

### eXplainable AI (XAI) methods

We calculated SHAP (SHapley Additive exPlanations)(73) values to quantify the individual feature contributions in predicting protein families using best-performing model

(HistGradientBoostingClassifier). SHAP values were extracted from the test set using the trained model. We then generated SHAP summary plots using the summary plot function from the SHAP library(20). To create SHAP bar plots, we calculated the total absolute SHAP values for each feature and determined the effect direction by computing the Pearson correlation between SHAP values and the model’s predicted probabilities

### Statistical analysis

Statistical Power tests(74) were performed using the TTestIndPower from the statsmodels package in Python to compute the Cohen’s d effect sizes for comparisons between groups (effect size = 0.8, alpha error = 0.05, power = 0.8). The Wilcoxon(75) test was used to compare the means and medians between feature sets and models. We ran Tukey’s test(76) pairwise comparisons among F1 scores to identify feature sets that significantly contribute to model performance. We applied p≤0.05 as statistical significance threshold across all statistical tests.

## SUPPLEMENTARY FIGURE LEGENDS

**Fig. S1. Principal component analysis of protein families and GO terms.**

A) PCA plot illustrating the variance captured by the first three principal components across different protein families. B) PCA plot depicting the variance explained by the first three principal components across Gene Ontology (GO) terms.

**Fig. S2. Global feature importance in Protein Family Classification.** Mean absolute SHAP values were computed for each protein family, representing the overall impact of features on each protein family. The direction of the impact was computed based on the correlation between the SHAP values and the likelihood of the predicted class: red for positive correlation, blue for negative correlation. Subplots (A-G) correspond to distinct protein families.

## SUPPLEMENTARY TABLE LEGENDS

**Supplementary Table 1**: Protein physicochemical features based on chemical properties of amino acid side chains (CPAASC).

**Supplementary Table 2**: Protein sequence composition features.

**Supplementary Table 3**: Distribution of selected protein families.

**Supplementary Table 4:** Distribution of selected GO terms.

**Supplemental Table 5.** MI Scores for protein families and GO terms.

**Supplemental Table 6.** Wilcoxon p-values for pairwise comparisons between protein families.

**Supplemental Table 7.** a) Feature Set Performance: A comprehensive ranking of all evaluated feature sets across all tested algorithms, sorted by F1-score. b) Algorithm Performance Overview: A ranking of algorithms based on their mean F1-score across all feature sets, offering insights into overall algorithm efficacy. c) Top-Performing Algorithms by Feature Set: For each feature set, the five best-performing algorithms are highlighted, facilitating the identification of optimal algorithm-feature set combinations. d) Algorithm Characteristics and Performance: A detailed ranking of algorithms, including descriptions of their underlying principles, classification strategies, and performance metrics.

**Supplementary Table 8:** Tukey’s Test for Feature Significance Based on F1-Score. This table presents the results of Tukey’s post-hoc test, assessing differences and statistical significance among features based on their F1-scores.

